# Neuroline: an integrated modular pipeline for FAIR multimodal neurophysiology data processing

**DOI:** 10.64898/2026.10.23.761011

**Authors:** Andreas Knoben, Andreas Franzke, Sietse Kiewiet, Mathis Bassler, Cyriel M.A. Pennartz, Conrado A. Bosman, Umberto Olcese

## Abstract

The adoption of FAIR principles for code and data sharing in systems neuroscience is hindered by the size and diversity of datasets, the multitude of data formats and the complexity of analysis workflows. These challenges have led to the development of various data standards and modality-specific processing and analysis tools. However, it remains difficult to integrate the diverse processed data streams originating from state-of-the-art neurophysiology experiments into a common format, which is crucial for cross-modal analyses and open, FAIR sharing of data. To address this, we developed Neuroline, a Python package for processing neurophysiology data (electrophysiology, optical imaging, and behaviour) that combines multimodal integration, modularity, and standardised output within a single pipeline. It leverages established packages and consolidates the resulting processed data into the standardised Neurodata Without Borders (NWB) format. Neuroline provides a unified interface while preserving flexibility and extensibility. Here we describe the implementation and challenges of a pipeline capable of integrating different types of neuroscientific data and packages. Finally, we address how Neuroline contributes to the sustainability and accessibility of neuroscience data and software.

## Introduction

Many recent studies in systems neuroscience rely on the integration of multiple complementary data streams, combining high-density electrophysiology, calcium imaging, and behavioural measurements such as operant responses and facial motion or pupil size extracted from video recordings^1,2^. Such multimodal experiments provide new opportunities to investigate questions related - for example - to neural representations of stimuli, motor movements, or context across brain regions and behavioural states^3–6^, to the way the brain implements cognitive functions^7–10^, and to how neuropsychiatric disorders affect circuit-level neural processing^11–13^. However, they also generate complex and heterogeneous datasets that are acquired, processed, and stored using a wide variety of instruments, software packages, and data formats, making data integration, reproducibility, and sharing increasingly challenging.

Practical implementation of large multimodal datasets is limited by fragmentation in data formats and processing packages (Figure 1a). Acquisition software and processing or analysis tools each often use a few specific formats that are not necessarily directly compatible. This fragmented landscape of instruments, formats, and analysis packages is not only challenging for researchers who have to decide which tools to use^14–17^, it also introduces variance when other investigators using different software attempt to reproduce a set of results^18–21^.

**Figure 1.**
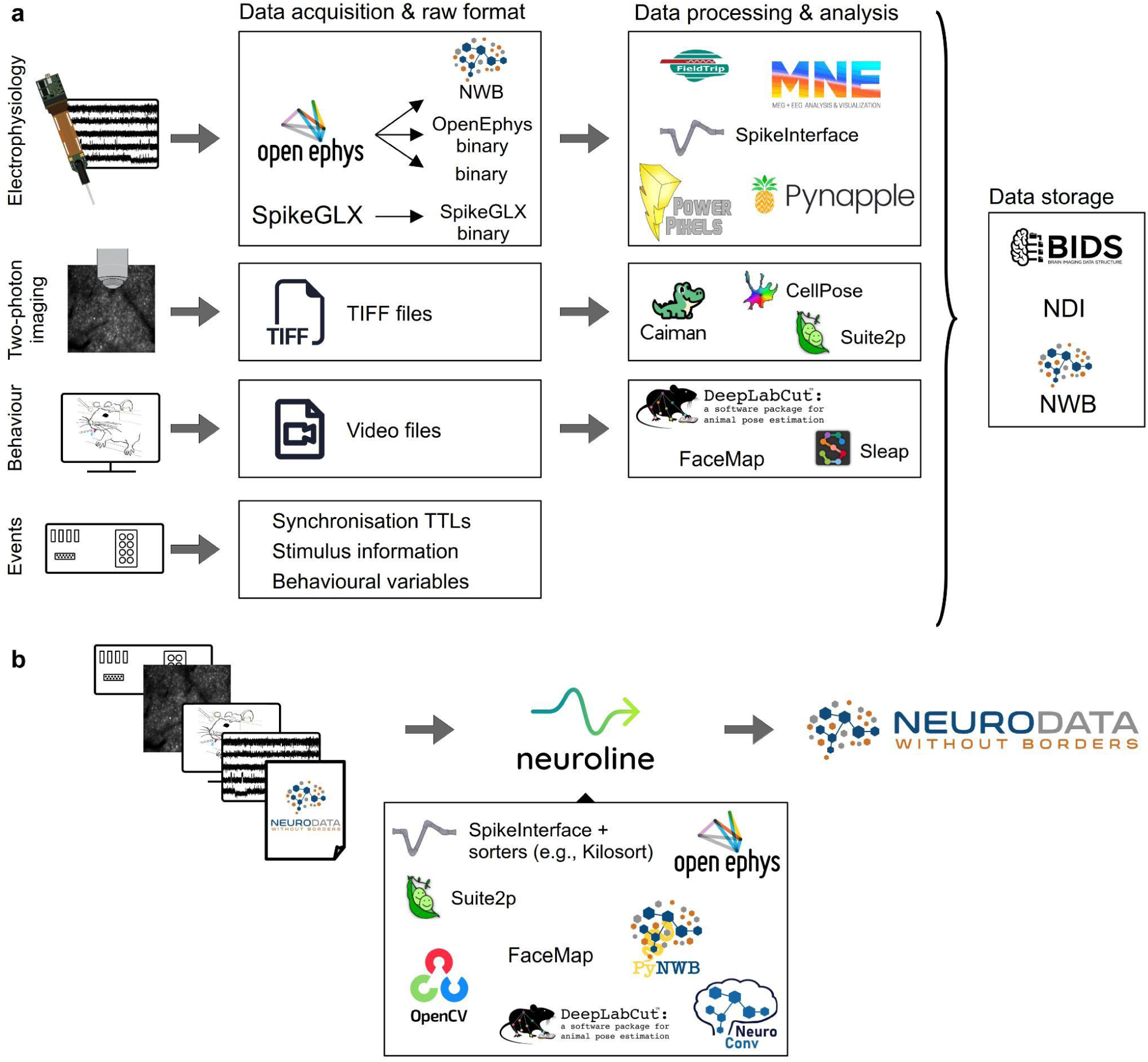
A plethora of choices present themselves at each stage of experimental data collection, processing and analysis in neurophysiology, which Neuroline simplifies. a) Each modality of data encountered in neurophysiology experiments, such as electrophysiological recordings, two-photon imaging, and animal behaviour, can be stored and processed using a large variety of tools and formats. At each stage of the pipeline (data acquisition, processing, and storage), the researcher has to decide which software package to use. Furthermore, data must be passed between different tools that each use their own set of preferred formats. Finally, the processed data must be combined into a standard data format to allow for easy sharing. This results in a large number of possible processing pathways. b) Neuroline aims to streamline this process by taking each type of data and processing it with specialised tools while providing an accessible and unified interface. This greatly reduces the number of choices a researcher is presented with and facilitates the process of passing data between packages and formats.

Such fragmentation also makes it harder to integrate all processed data into a single, standard output data format, the need for which is highlighted by the increasing importance of FAIR (Findable, Accessible, Interoperable and Reusable) principles^14,22,23^. These principles entail that research groups make their data and resources available in such a way that they can be easily found, re-used and integrated. For laboratories, this necessitates harmonising their data into a standard format while ensuring relationships and interdependencies between different parts are maintained. However, due to the vast variety in available pipelines and tools, such standardisation is not trivial and requires significant effort from the researcher. The resulting lack of interoperability often causes laboratories to maintain their own software ecosystem, duplicating efforts and limiting adoption of common, standardised tools^22^.

To address these problems, we developed Neuroline, a modular software pipeline written in Python for processing and exporting the diverse types of data collected in state-of-the-art *in vivo* systems neuroscience experiments. The package integrates existing processing tools and formats primarily for animal neurophysiology and behaviour, such as extracellular electrophysiology and two-photon imaging. Its flexible, modular architecture allows for straightforward extension to other subfields, including human electrophysiology recordings such as electroencephalography (EEG), magnetoencephalography (MEG), electrocorticography (ECoG) or stereo-electroencephalography (sEEG). Neuroline’s principal contribution is the combination of three features within a unified framework: multimodal integration, a modular architecture, and standardised output.

First, it integrates widely adopted and actively maintained (pre-)processing tools for multiple experimental data modalities. This reduces the number of choices researchers must make regarding analysis packages and avoids introducing a new analysis package by re-using existing neuroscientific software. Crucially, this integration enables multimodal processing, in contrast to many existing pipelines focused on a particular modality (e.g. ^24–28^). Neuroline orchestrates these tools and combines their results into a standard output format (Figure 1b).

Second, Neuroline provides a modular framework with a unified interface in which users can select which modules to run and specify their parameters using a high-level scripting syntax. Although the package provides a pre-selected set of dependencies, its existing scaffolding allows new data modalities and processing tools to be added and integrated seamlessly into the pipeline.

Third, the resulting processed data are written into the standard Neurodata Without Borders (NWB) format^29^. This supports the organisation and sharing of processed data in a way consistent with FAIR principles. Moreover, using a standard data format facilitates compatibility between Neuroline’s output and downstream analysis tools^30–32^. Together, these aspects allow Neuroline to serve as a standardised pipeline for multimodal system neuroscience, while remaining agnostic to the specific experimental modalities used.

Importantly, despite promises of standardisation and interoperability brought by existing tools, development of Neuroline revealed that challenges remain towards implementing FAIR principles, particularly in passing data between packages, loading data from different sources and handling diverse experimental setups, and properly combining all processed data into the output data format. Neuroline resolves these issues by providing flexible data loading interfaces and handling formatting when piping data between packages where necessary, ensuring data are complete and correctly formatted.

This manuscript provides an overview of Neuroline. First, the structure of the pipeline and a typical workflow are described, followed by a practical use case. Finally, the challenges encountered during development, and how they have been addressed in Neuroline, are discussed.

## Results

### Software overview

Neuroline is a modular software package that integrates state-of-the-art analysis tools for extracellular electrophysiology, optical imaging, and animal behaviour into a unified workflow (Figure 1b). The pipeline can pre-process raw data, handle processed data, and export the results into the Neurodata Without Borders^29^ (NWB) format, an open-source standard for neuroscientific data storage endorsed by the International Neuroinformatics Coordinating Facility, INCF^14^. The NWB format is supported by an ecosystem of storage, conversion, and analysis tools in different programming languages^31^. The use of this standard streamlines data sharing within and outside of the lab and contributes to the *findability* and *accessibility* principles of FAIR science by providing a documented, machine-readable schema.

The package, which is written in Python, currently accepts raw binary or NWB data from the OpenEphys electrophysiology acquisition software^33^; processed electrophysiological data, including spike-sorted data or local field potentials; two-photon imaging data as Tag Image File Format (TIFF) image stacks or Suite2p^34^ output; behavioural videos, such as orofacial recordings of an animal; and behaviour and stimulus information, including event timings coded by transistor-transistor-logic (TTL) events.

Neuroline is entirely modular, allowing the user to run only the modules they need. This modularity also provides flexibility for the pipeline to be extended relatively easily to new types of data. Each module takes the relevant data and processes it with the steps and options the user provides, with effort put into ensuring reasonable parameter values are provided by default, such as best practices in the field or recommendations by underlying software packages. The processed data can then be exported into a new or existing NWB file.

### Software modules

Neuroline comprises three top-level modules (Figure 2). The *instances* module contains objects representing a particular part of the pipeline, such as a device or recording. These objects tie an experimental recording session together, provide the necessary metadata, hold objects that point to the data, and execute and orchestrate the pipeline. *Processing* consists of collections or submodules containing modality-specific processing functions. Each submodule shares a common structure and provides an accessible and unified interface to existing packages. Finally, the top-level *utils* module consists of general utility functions, mostly used internally by the pipeline. Crucially, these modules are not fixed components. For instance, adding a new data type or recording modality only requires implementing the necessary *instances* and *processing* modules. As long as these modules conform to the existing structure, they integrate automatically with the rest of the pipeline.

**Figure 2.**
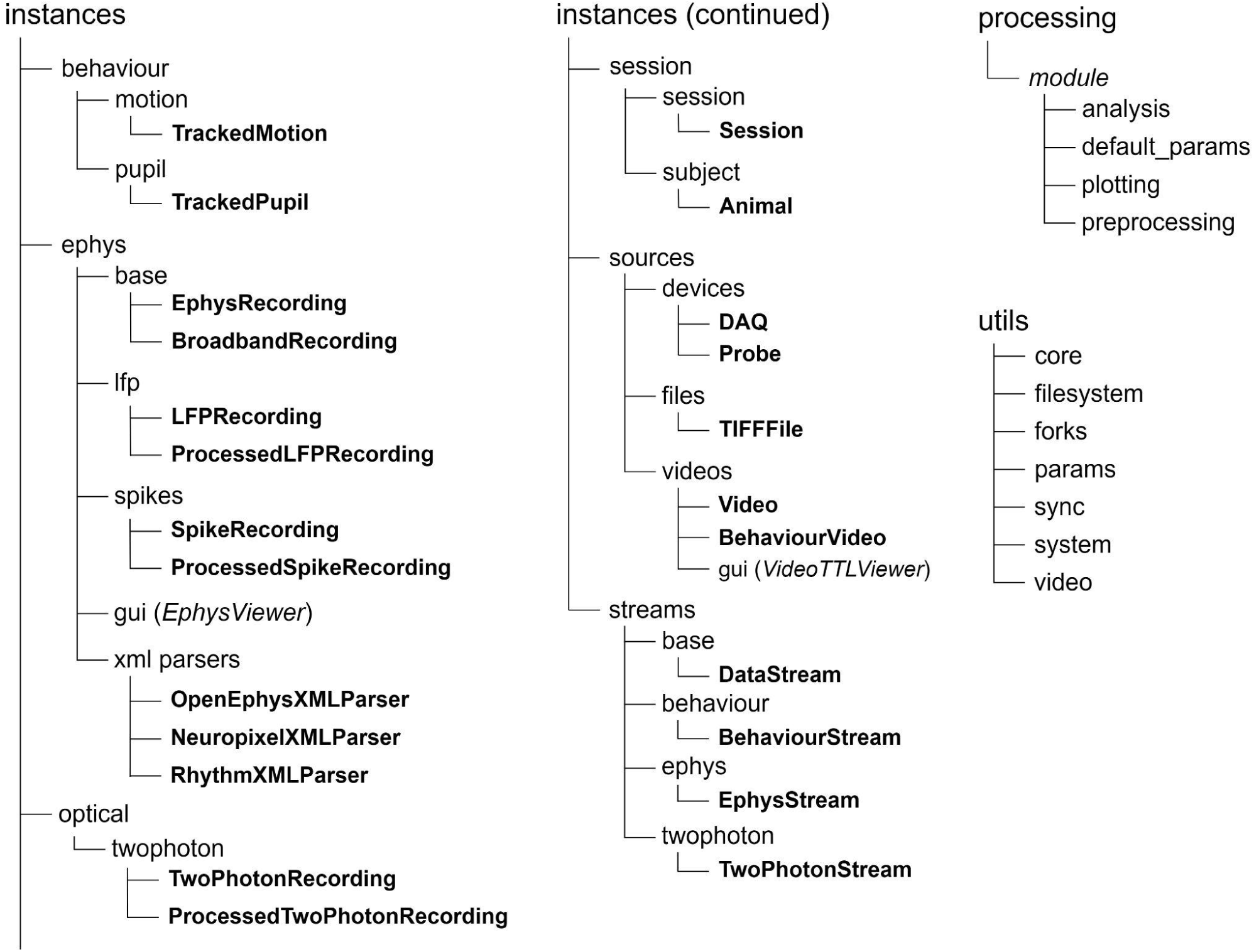
Technical structure of the Neuroline package. The package has three top-level modules, namely *instances*, *processing*, and *utils*. The *instances* module contains objects that represent session information (in *session*), data sources (in *sources*), data streams (in *streams*), or a particular type of data (in *behaviour*, *ephys*, and *optical*). The *xml parsers* module contains classes that parse an OpenEphys XML (Extensible Markup Language) file to obtain session and recording information. Some modules, such as *ephys*, also contain a graphical user interface (*gui*) to validate processing steps. The instances themselves are made bold in the figure. The *processing* module contains data processing ‘submodules’, collections of functions for a specific modality that can be processed with Neuroline (currently *lfp*, *spikes*, *twophoton*, *motion*, *pupil*). Each processing submodule contains a set of default parameters, preprocessing functions, and visualisation functions. Sometimes, a processing module may also comprise analysis functionality, i.e., functions that can help the user in later analysis stages. Finally, the *utils* module contains utility functions, most of which are used by the pipeline internally.

Instances are structured hierarchically within the package. At the top of this hierarchy stands the *Session* class, which holds information about the experiment, subject, and registered data streams represented as *DataStream* objects. These stream objects form the “second layer” and handle a specific type of data, such as an *EphysStream* for an electrophysiological recording. The streams manage the necessary information for a particular type of data and hold one or more data sources. These sources can be synchronised when a data acquisition device (DAQ) associated with the stream provides timing information. A DAQ object may be associated with multiple streams, in which case Neuroline assumes the streams share a timing reference and that their relationship can be established through recorded synchronisation events such as TTL pulses. These events are used to assign timestamps and align streams, although the exact implementation depends on the type of stream and the available information. For example, timestamps of electrophysiology probe data are related to a reference clock through a linear fit, defined by a scaling factor and offset. Data points can be assigned timestamps from digital or analog TTL pulses. Alternatively, timestamps can be inferred between synchronisation events, such as start and end TTLs.

In a typical use case, an experiment may include an *EphysStream* and a *BehaviourStream* referring to a video recording of an animal’s behaviour. The *DAQ* instance from the *EphysStream*, representing a data acquisition device containing timestamps and TTL events, can also be associated with the *BehaviourStream*. Timestamps can then be assigned to the video by using the TTL events corresponding to video frame acquisition, so that the two streams can be aligned in the final NWB file.

Data sources constitute the “third level” of instances. They can be a *Device*, such as a *Probe* instance for electrophysiological recording probes, a *File*, for example, a *TIFFFile* from a two-photon imaging experiment, or a *Video*, such as a *BehaviourVideo* referencing an orofacial motion capture. Data source instances hold metadata about the recording as well as pointers to the actual data on the disk. Furthermore, they have methods relevant to each device, file, or video, such as extracting a subset of channels from a probe or loading events from a data acquisition system.

The fourth and lowest level in the hierarchy consists of “data-representing” instances, which are derived from a data source. For example, a *TIFFFile* may yield a *TwoPhotonRecording*, or a *BehaviourVideo* may yield an instance of *TrackedPupil*. Data-representing instances have (pre-)processing methods which call on the *processing* submodules to execute various steps in a pipeline. These processing modules use existing packages as much as possible to produce a “processed data” object, e.g. calling *preprocess()* on an *LFPRecording* results in a *ProcessedLFPRecording*. The executed (pre-)processing steps, their order, and their options can be determined by the user. Ultimately, the processed data can be added to an NWB file from the processed-data-representing instances.

While this “layered” approach may seem complex, it provides the flexibility necessary to process an arbitrary number of different types of data from various sources, while still being able to manage the experiment as a whole. The latter is important for synchronisation and integration of the different data streams. Furthermore, this flexibility in the pipeline makes it extendable. Adding a modality only requires implementing some or all of the following components: a data stream, data source, data-representing object, or processing submodule. Because of the modular structure, these can be added without disrupting the existing pipeline. Moreover, the existing code provides scaffolding in the form of abstract and base classes that aid in the implementation of inheriting classes.

For example, implementing functionality to load and process widefield calcium imaging data from binary files would be achieved by a *WidefieldStream* inheriting *DataStream*, a *BinaryFile* source class inheriting *File*, a *WidefieldRecording* class, and a *widefield* processing module. These latter two can follow the structure of other optical imaging classes, such as *TwoPhotonRecording*.

### Workflow

The hierarchical structure of the package can be seen clearly in its general workflow (Figure 3). It starts with the creation of a *Session* instance. This instance requires various pieces of information about the experiment, such as the session’s root path, names of the experimental paradigms run during the recording, and subject information, e.g. an *Animal* instance providing the subject’s age, sex, genotype, and other relevant properties. After the *Session* has been initialised, users can add as many data streams as necessary. When a data stream is constructed (e.g., *EphysStream*), it automatically constructs the relevant data source objects (e.g., *Probe*). In turn, these data sources create the corresponding data-representing objects (e.g., *BroadbandRecording*). Each parent object maintains references to its child objects, allowing users to access them through the object hierarchy.

**Figure 3.**
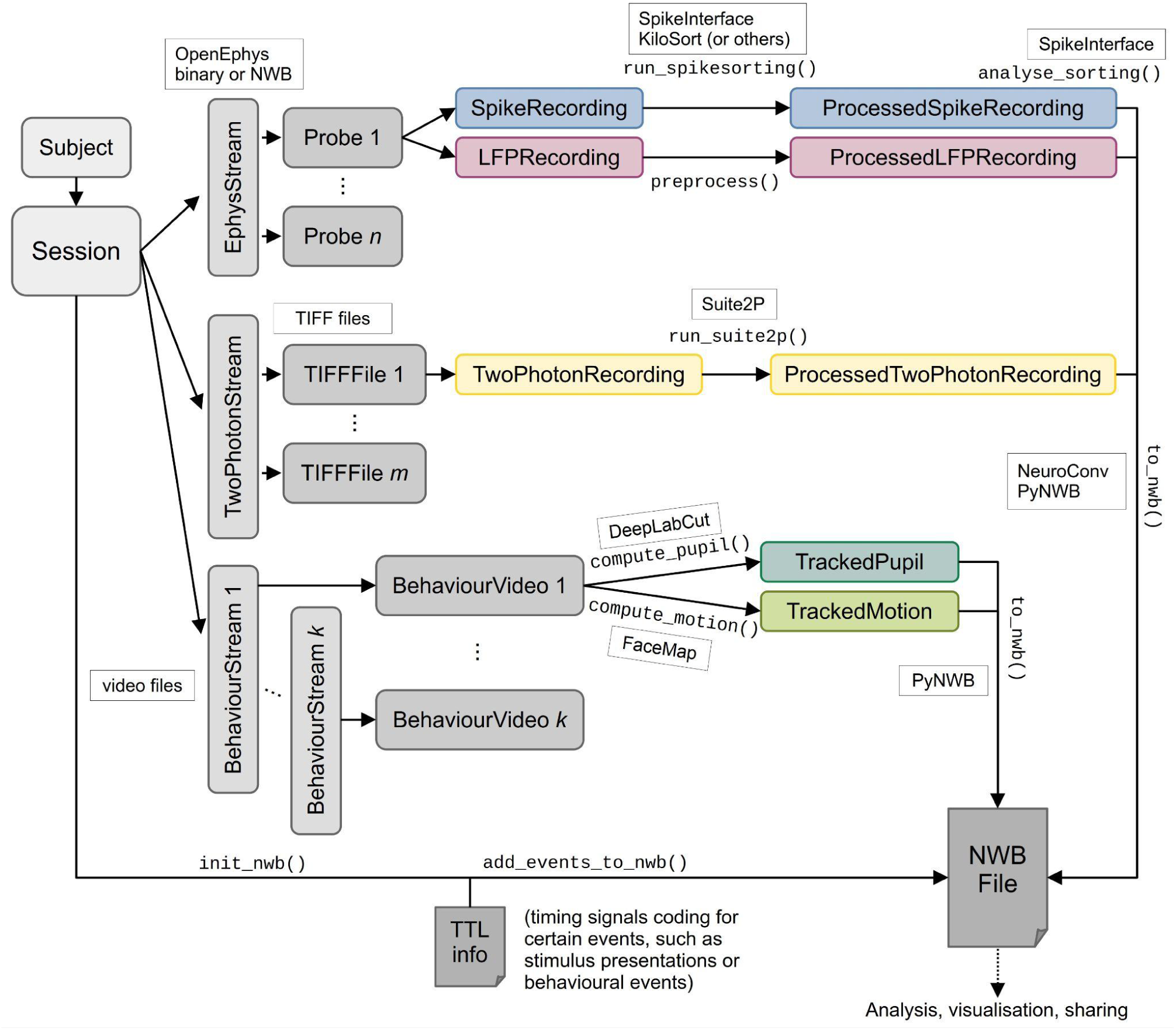
Visualisation of the workflow of Neuroline. The graph depicts the workflow for a recording of electrophysiological data using *n* probes, two-photon data stored in *m* TIFF files, and *k* behavioural videos. The rounded rectangles represent objects or instances; the arrows represent what instances can be produced by a particular object, with the adjacent monospace text representing the corresponding method; the smaller, square rectangles show what external files or packages are used in a particular instance or method. A *Session* object is created, containing information about the subject (e.g. *Animal*). In this example, it holds the following data streams: an *EphysStream*, which contains the *Probes*, representing silicon probes for extracellular electrophysiology, and their associated *SpikeRecording* and *LFPRecording*; a *TwoPhotonStream*, containing references to the TIFF files (*TIFFFile*) and their associated *TwoPhotonRecording*; multiple instances of *BehaviourStream*, which link to the associated videos (*BehaviourVideo*). The spiking data and LFPs can be preprocessed into *ProcessedSpikeRecording* and *ProcessedLFPRecording* objects respectively; Suite2p^34^ can be run on two-photon data for cell detection and computation of fluorescence levels over time to get a *ProcessedTwoPhotonRecording*; pupil tracking data (*TrackedPupil*) and orofacial motion (*TrackedMotion*) can be extracted from the videos. After initialising the NWB file by calling *init_nwb()* on the *Session* instances, all processed data can be added to the NWB file by calling *to_nwb()* on each processed data object. Furthermore, TTL events and stimulus information can be added using *add_events_to_nwb()* from the *Session* instance.

After this initialisation is complete, the user can run different preprocessing and processing steps on each of the derived data-representing instances, for example, synchronising the data to a common temporal reference, running spike sorting and sorting analysis on spiking data, preprocessing the LFP data, extracting cell fluorescence from two-photon data, or computing motion energy from a video (see Figure 4 for example code). After the processing steps are run on a data instance, a processed data object is returned, containing references to the processed data on disk.

**Figure 4.**
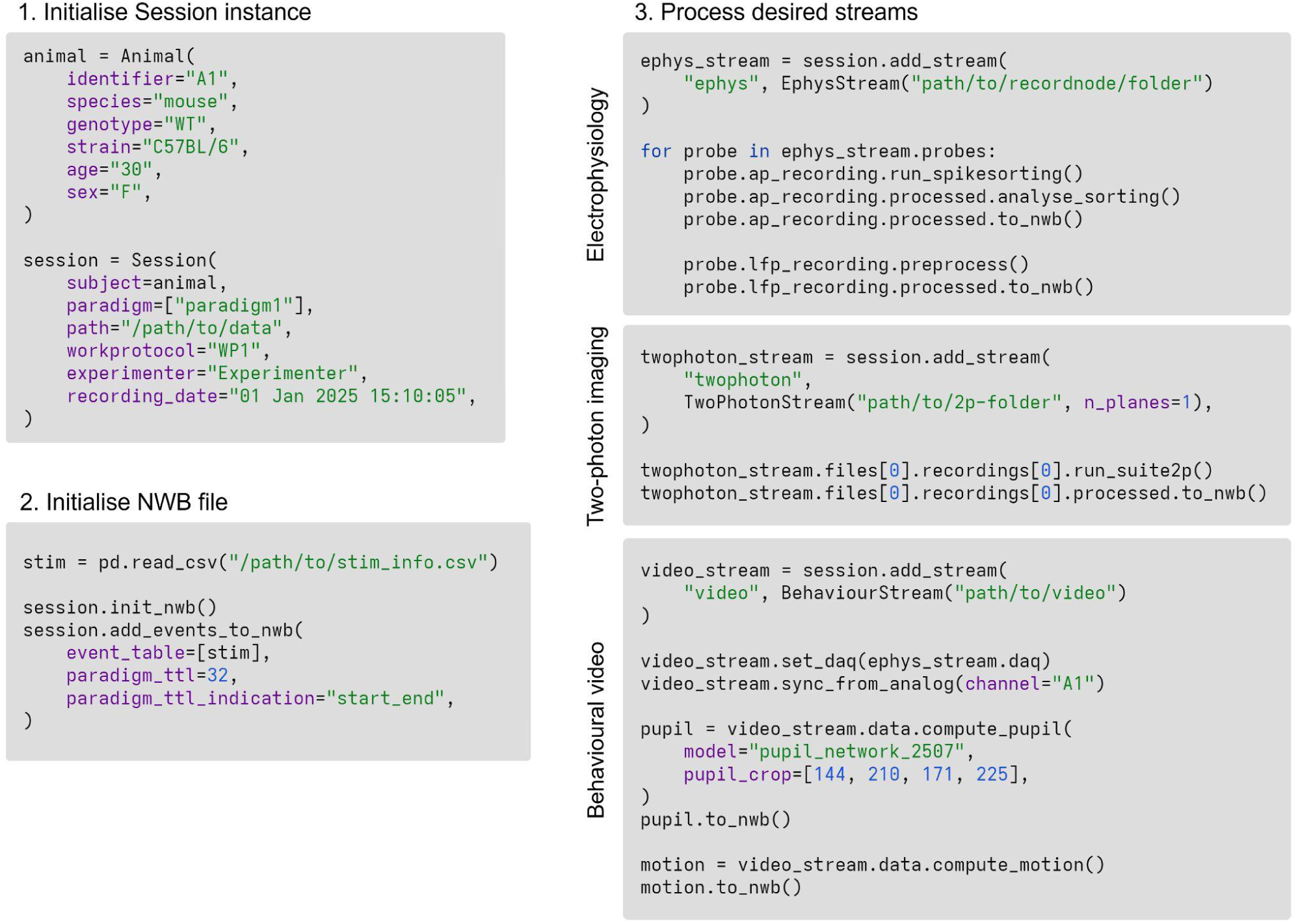
Example Python scripts demonstrating the interface to Neuroline. The first step is initialising a *Session* instance, which requires a subject, e.g., an *Animal*. Next, an NWB file can be initialised with stimulus or event information. To process specific types of data, their corresponding streams must be constructed and added to the *Session*. Then, for electrophysiological data, spike sorting and LFP preprocessing can be run on each probe, and the resulting spikes and LFPs can be added to the NWB file. For two-photon data, after the stream is created, Suite2p is run on the recording and its output is added to the NWB file. For the behavioural video data, first timestamps are assigned by relating the data acquisition device (DAQ) from the electrophysiology stream and synchronising using the channel that received the camera’s pulses. Finally, pupil tracking and facial motion extraction are run on the video and subsequently appended to the NWB file.

Finally, to get a standardised output, an NWB file must first be initialised from a *Session* instance. Because the NWB file is generated from the information available in this object, the required metadata is inserted into the file automatically. Such metadata includes, for example, experimental information, such as subject, the paradigms that were run and information about the presented (sensory) stimuli, but also modality-specific notes such as details of equipment or processing steps. Next, the user can add stimulus and event information, consisting of tables with properties of presented stimuli and events and their associated TTLs. Finally, the processed data and associated metadata can be added to the NWB file using Neuroline’s built-in export functionality, without requiring the user to manually construct each NWB container. The processing steps and package versions used in the environment are included to preserve the provenance of the analysis and facilitate its later interpretation and reuse.

### Installation and dependencies

Neuroline is open source and available on GitLab (https://www.gitlab.com/csnlab/neuroline) and the Python Package Index (PyPI). User guides and documentation are also available on GitLab (https://csnlab.gitlab.io/neuroline). To accommodate differences in hardware and software configurations, users must first create a dedicated Python environment using the conda package manager^35^. Within this environment, PyTorch^36^, a Python library for deep learning on which several packages integrated in Neuroline depend, is installed before Neuroline itself. The documentation provides instructions for different computer systems and configurations.

### Use case: Determining the contribution of sensory, movement and arousal related variables for *in vivo* electrophysiology

To demonstrate the pipeline’s practical applicability, we applied it to a multimodal neural dataset comprising *in vivo* extracellular electrophysiological data collected while an animal performed a multisensory detection task (Figure 5A) ^5,37,38^. Specifically, the mouse was trained to indicate whether a change in sensory stimulation occurred in the auditory domain (i.e., a frequency change) or visual domain (i.e., an orientation change) by licking to the right or left lick spout, respectively. Correct trials were rewarded with a small amount of milk. A key question in experiments such as this one is how experimenter-controlled task events, self-generated behavioural events, and arousal state contribute over time to single-trial neuronal firing rates.

**Figure 5.**
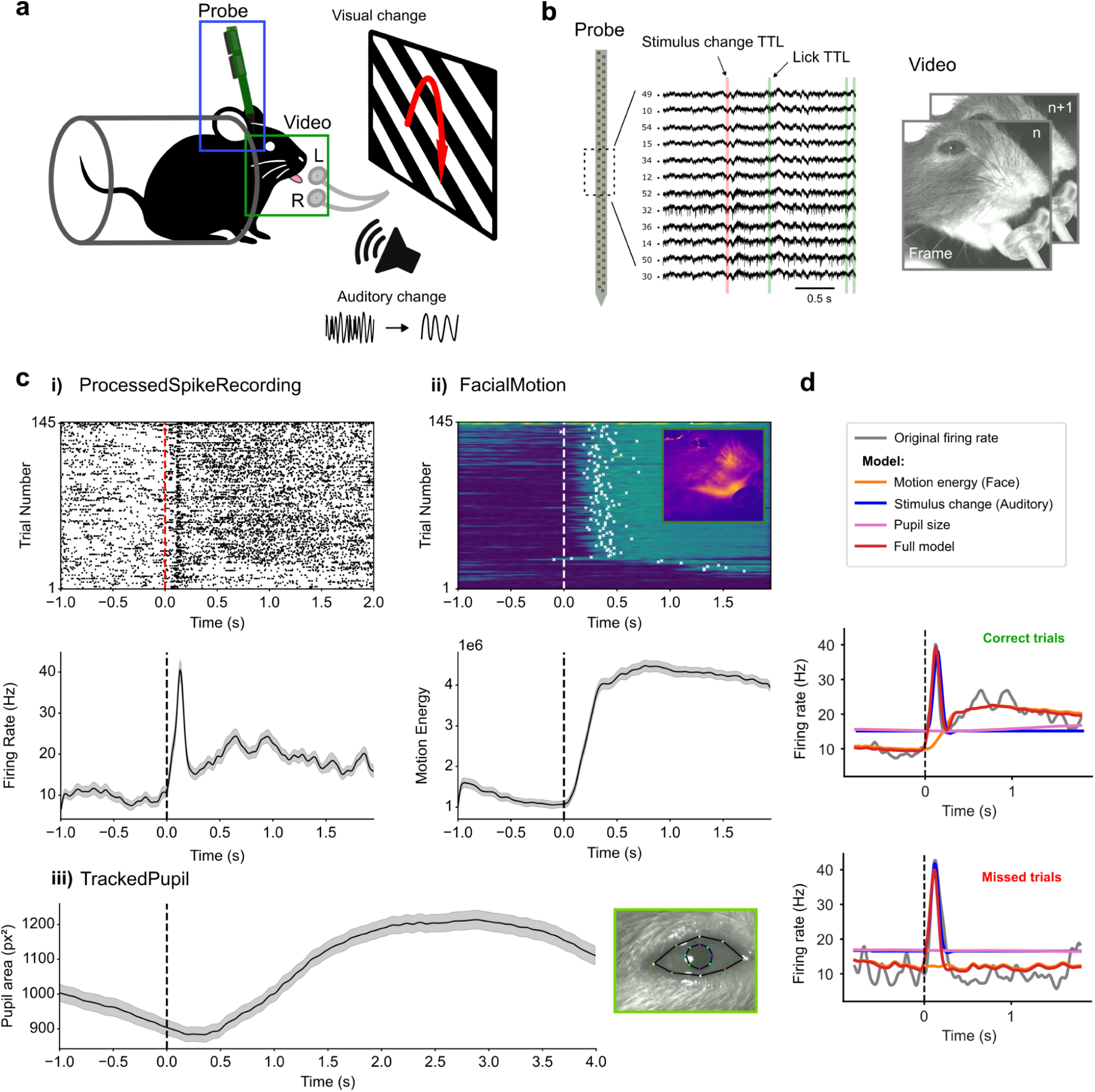
Use case: Determining the contribution of sensory, movement and arousal related variables during in vivo electrophysiology. **a)** Schematic image of the experimental setup: a head-fixed mouse is continuously presented with auditory and visual input. The mouse was trained to indicate whenever a change occurs in either sensory stream (frequency change [Hz] vs visual change [degree]), and could obtain a small milk reward by licking to the correct side. b) Visualisation of the raw data streams. Left: extracellular electrophysiological data were collected in the auditory cortex using a NeuroNexus 64-channel silicon probe. Presented are 12 channels with overlaid TTL pulses, indicating the onset of an auditory stimulus change in red, and multiple lick TTLs highlighting the times at which the animal licked in green. Right: a video of the face is captured at 100 frames per second (fps). c) Visualisation of the processed data streams. i) *ProcessedSpikeRecording*. Top: Spike raster plot for an example single neuron, plotted for all auditory trials and aligned to auditory stimulus change. Trials are sorted by motion energy magnitude (*see* ii). Bottom: peri-event time histogram (PETH) of the same neuron and trials shown in the top plot. Data are presented as the mean ± standard error of the mean (sem) **ii)** *FacialMotion*. Facial motion energy (ME) per trial, sorted by ME magnitude (mean over the whole trial). The first registered lick is visualised as a white cross overlaid on top of the motion energy data. Bottom: mean ME for the trials shown in the top plot (mean ± sem). **iii)** *TrackedPupil*. Left: mean pupil area (pixels squared) during the same auditory trials shown in i-ii (mean ± sem). Right: pupil with “key points” as predicted by the DeepLabCut model visualised for an example frame. **d)** Single neuron firing rate was modelled using a Generalized Linear Model (GLM) including only ME predictors (orange), including only auditory stimulus predictors (blue), using only pupil size predictors (pink) and a “full” model, including all predictors (red). For visualisation purposes, a small offset was applied to the predicted firing rate of the auditory-only model.

To address this question, complementary raw data streams were acquired, including 64-channel silicon probe recordings from primary auditory cortex, behavioural video recordings of the animal, and multiple behaviour- and stimulus-related TTL events (Figure 5B, see *Methods*). The raw data streams were processed with Neuroline using user-selected pre- and post-processing parameters. The resulting processed objects were a *ProcessedSpikeRecording*, a *FacialMotion*, and a *TrackedPupil* instance. Once the raw data processing had been performed, the streams were automatically synchronised and saved in a standardised NWB file. The individually processed data streams were then visualised using existing packages that work with time series data and support the NWB file format, such as Pynapple^32^ (Figure 5C i-iii).

A particularly well-suited approach to examine the relative contributions of task-related, movement-related, and arousal-related variables to neural activity (Figure 5C i) is to model the firing rate using a Generalised Linear Model (GLM), here implemented with the NeMoS package^30^ in combination with Pynapple^32^. Because all individual data streams are temporally aligned within a common reference frame, behavioural variables and task-related events could be used directly as predictors in the model, providing an estimation of the contribution of each predictor stream to single-neuron firing rates (Figure 5D). The initial peak in neural activity could be largely predicted by task-related behavioural events (blue line: presence of auditory stimulus change), while movement-related variables mainly contributed later in time, relative to stimulus change (orange line: facial motion energy) and only during ‘correct’ trials, indicating that the recorded neural activity was indeed jointly driven by both stimulus and movement variables. Pupil size, a proxy for arousal^39^, showed only a small predictive contribution late in the post-stimulus window (Figure 5D, ‘correct’ trials).

This example illustrates how Neuroline streamlines the process of going from raw multimodal data acquisition to quantitative analysis within a couple of steps. Data acquired in heterogeneous formats (e.g. binary, MP4) are processed, synchronised, and standardised in a common data output format, ready for downstream analysis tools such as Pynapple^32^ and NeMoS^30^. In this way, Neuroline can facilitate the reusability and reproducibility of analysis procedures within and across labs.

To further demonstrate the functionality of our package, we also applied it to a two-photon recording of spontaneous activity from primary visual cortex (V1) and lateromedial secondary visual cortex (LM), performed in a mouse expressing the GCaMP6f calcium indicator (see Bassler et al., 2026^40^ for details; Supplementary Figure 1). The recording TIFF file was loaded into Neuroline to yield a *TwoPhotonStream*. Suite2p^34^ was called by Neuroline to detect cells, extract their fluorescence signals over time, and infer spikes by deconvolution of the fluorescence traces. This resulted in a *ProcessedTwoPhotonRecording* instance, from which the segmentation images and fluorescence traces were added to the NWB file.

## Discussion

Neuroline addresses substantial challenges that remain in the synchronisation, documentation, and integration of outputs in multimodal neurophysiology. Large, heterogeneous datasets are often generated by multiple recording systems and processed by specialised tools, making it difficult to preserve relationships between streams and represent them in a common format. These requirements are particularly important in the context of FAIR and open science, which emphasise making data findable, accessible, interoperable, and reusable^14,22,23^. Constructing such a pipeline from scratch not only demands substantial development work but also presents researchers with a multitude of choices regarding packages and data formats. Neuroline addresses this integration problem by providing a modular framework that organises existing packages and their outputs. It provides an interface through which different modalities of data can be processed, synchronised, and exported to the NWB standard.

Neuroline is designed around three principal features: 1) multimodal data processing and integration; 2) a modular architecture with a unified, high-level interface; and 3) export of data streams in NWB format. In the following sections, these features and the limitations associated with these aspects will be discussed. We also consider challenges encountered during development that are related to integrating different types of data.

### Capabilities and limitations

#### Multimodal data processing and integration

Although existing packages are very suitable to process individual data modalities (e.g., DeepLabCut^41^, FaceMap^42^, MNE-Python^43^, SpikeInterface^24^, Suite2p^34^, SyNCoPy^44^), combining the resulting data remains challenging. Different data streams have to be synchronised and integrated so they can be related to each other. Moreover, for the purpose of sharing, it is convenient to store different types of data together. By combining processing packages for different modalities into a unified workflow, Neuroline facilitates synchronisation and joint storage of multimodal data while harnessing the power of specialised tools and their community support. This integration reduces the effort required to combine and align data and facilitates subsequent cross-modal analyses, such as prediction of neural activity from behaviour. Furthermore, as the pipeline runs, it can produce visualisations to validate processing steps and synchronisation.

#### Modularity without complexity

Neuroline integrates modalities and packages while preserving modularity: users can choose which modules are run and determine their order, and they can implement their own modules when needed. New data modalities can be implemented by building upon the existing structure of the package without interfering with other modules. Importantly, this flexibility is exposed through a user-facing API designed to reduce the amount of workflow and data-format engineering required from researchers, as Neuroline can be operated using relatively simple Python function calls.

Although there are software packages that offer a similarly flexible pipeline allowing for integration of different data sources^27,45–47^, they are typically more difficult to adopt for users without a strong background in software engineering. For example, DataJoint-based pipelines require some understanding of relational databases^27,47^. Rule-based systems, which use a custom syntax to describe data transformations, or git-based systems, which use the version control system git for provenance tracking and hierarchical processing, introduce their own conventions to orchestrate workflows^45,46^. In addition, some of these pipeline builders still require the user to implement the underlying processing and formatting steps^45,46^, meaning they may not fully address the issues of package selection, processing variability, and data integration.

In contrast, Neuroline manages data streams, the transfer of data between packages, and the combination of results into a standardised output, accessible through a simple Python API. However, such an abstraction also entails limitations. Neuroline provides less fine-grained control over the included tools and their configuration compared to lower-level workflow systems. Additionally, database-backed, rule-based, or git-based alternatives may offer more powerful provenance tracking. Neuroline also does not replace systems backed by databases or large-scale distributed computing frameworks.

#### Standardised output

Not all packages are able to collect the final output in a standardised format natively (e.g. ^24,41,42,45–47^). Neuroline generates NWB files, a format endorsed by the INCF (International Neuroinformatics Coordinating Facility)^14^, containing the processed data and associated metadata. Although NWB has been chosen as the pipeline’s output format, the modularity of Neuroline allows for integration of other formats. Notably, we aimed to leverage existing infrastructure to support FAIR practices: NWB’s constraints on file structure aid *findability* and *accessibility*; *interoperability* is promoted by using existing packages and offering output compatible with the NWB ecosystem and other (downstream) packages across programming languages that are able to read NWB data; *reusability* is supported by providing a modular, flexible structure, API documentation, and (abstract) base classes to build upon.

However, generating an NWB file does not by itself guarantee full FAIR compliance. Neuroline provides a supporting framework to organise data and metadata, but the quality and completeness of the resulting output depends on the information supplied by the user and on the adopted documentation and sharing practices. Neuroline itself does not currently enforce any strict checks on metadata completeness or quality and still requires user oversight and input to generate a complete and adequately documented representation of the data and the processing steps.

#### Challenges encountered when integrating systems neuroscience software

During the development of Neuroline, we encountered a number of issues, particularly when integrating different packages. These problems highlight outstanding challenges in standardisation, interoperability, and sustainability of systems neuroscience software. Addressing these issues is important for Neuroline and other analysis tools to reliably integrate other packages. Although some of these issues are subject to change in the longer term, we now describe how Neuroline addresses them currently.

#### Package-format interoperability

The first key problem pertained to the interoperability between different data formats and analysis packages. For example, extracellular electrophysiology data acquired using OpenEphys can be stored in binary or NWB format. The choice of format influences the amount of information that must be supplemented manually to construct a correct and complete SpikeInterface^24^ recording object, which is the package we chose for preprocessing electrophysiological data of any provenance. However, even when the data come from the same acquisition software, different recording formats require distinct processing pathways. In a similar fashion, when using automated NWB conversion tools (e.g., NeuroConv^48^), which aim to automatically format and place data into an NWB file, not all data might be converted into the output. For example, when attempting to automatically write spike sorting output along with sorting quality metrics into an NWB file, some metrics necessary for further analysis, such as unit location, were not written into the file and had to be supplied manually. In these types of cases, Neuroline was designed to add missing data conforming to schema guidelines.

These kinds of problems make it more difficult to chain different tools together, requiring the researcher to invest extra time into programming a solution, e.g. supplying extra metadata or formatting the data differently, and verifying the results are correct. Neuroline addresses this directly by ensuring the data is appropriately structured and all necessary metadata is included when it is passed into a tool. This is done either by implementing existing tools that prepare data for inclusion into NWB such as NeuroConv^48^ or by placing the data in appropriate structures conforming to the NWB schema specification^29^. That way, powerful tools can be used together without placing extra burden on the end user.

#### Incomplete or inconsistent documentation

The second problem we faced during development was incomplete or inconsistent documentation of the packages we were integrating, particularly for niche functionality or for inspecting a function’s behaviour. Verifying the results of a processing step or understanding the inner workings of a function often required digging through the source code as documentation would not provide this information. Furthermore, documentation was frequently re-used across guides, tutorials, and API (application programming interface) references, with discrepancies between the information provided in each source. This made verification of the behaviour of a package more difficult, which hindered identification of sources of bugs, artefacts, or unexpected results. Providing usable, correct, and up-to-date documentation is an essential factor in package adoption and hence in reproducible and collaborative research^15,49,50^.

We have addressed this by adopting a two-part documentation consisting of high-level guides and a detailed API reference. The high-level guides allow end users to get started with the package relatively easily, while the API references provide users and developers with an in-depth understanding of available functions and their parameters. Care is put into ensuring the guides are kept updated with code changes, whereas the API references are generated automatically from the code and the comments therein. Furthermore, since Neuroline, including its documentation, is open source, the community can contribute to the guides and API text, strengthening verification and relevance.

#### Rigidity of format schema

A third issue we encountered concerned the storage of the processed data within the ontology of the standardised format selected for Neuroline, NWB. While such schema definitions are designed to ensure consistency between datasets^29^, their specificity can become restrictive when data deviate from the types of objects anticipated by the schema. In such cases, storage location and format become unclear. Different decisions may then introduce variability between datasets despite the use of the same standardised format. Other labs have reported similar issues when storing data that do not conform strictly to the assumptions of the NWB format^1,51^.

An example of this problem presented itself when storing FaceMap’s singular value decomposition (SVD) output. An SVD applied to a video yields a set of motion components, each of which is associated with a spatial mask and a time series. Although time series can generally be stored in an NWB *TimeSeries* object, the NWB schema does not specify a specific time-series container for this kind of motion as it does for other measurements, such as pupil tracking. More difficult, however, was the decision on how to store the motion masks. We therefore stored them in a *TimeSeries* object because this structure accommodates arrays of any dimensions. However, this is semantically incorrect as the masks represent spatial information rather than a time-varying signal. Hence, this is an example of a temporary but documented implementation until a more appropriate container is available.

Rather than presenting such decisions to the end user, Neuroline has implemented informed and documented mappings for cases in which such an “ontological gap” arises. This ensures that outputs are stored in the most suitable available NWB container with a consistent metadata representation. Neuroline is maintained to track and implement updates to the NWB schema, so mappings can be revised when a more appropriate container is provided.

#### Package sustainability & fragmentation

A more fundamental challenge was selecting the packages and formats on which to base the pipeline. A key consideration in such a selection is the expected sustainability of these tools so that they retain compatibility with other packages and new Python versions. This challenge is exacerbated by the existence of alternative tools for the same task. For example, there are several packages for electrophysiology data processing, including Elephant^52^, MNE-Python^43^, SpikeInterface^24^, and SyNCoPy^44^. While this diversity offers researchers flexibility, it can make it difficult to select and maintain a mutually compatible set of tools.

Neuroline aims to reduce the uncertainty surrounding these decisions by integrating well-supported, widely used packages. These packages have been chosen based on project activity, discussions with researchers, and available literature. Neuroline also manages package versions and dependencies, reducing the effort required from users to maintain a compatible software environment. The versions of the packages are stored with each data output to provide a transparent record supporting reproducibility. Updates to integrated packages are incorporated with the aim of minimising user-facing changes while leveraging updated functionality and bug fixes. A minimal test suite is used to validate the pipeline, test different types of inputs, and identify potentially breaking changes.

However, dependency management shifts part of the maintenance work to the Neuroline developers. It is therefore possible that Neuroline lags behind upstream releases when incompatibilities or interface changes require updates to Neuroline’s code. Additionally, the selection of implemented tools may reduce users’ immediate access to alternative packages, but the modular architecture allows for additional tools to be incorporated when needed.

#### Software compatibility

Finally, a core aspect for improving interoperability between different packages is designing software that is compatible with tools that precede or follow it in an analysis workflow. Recent examples are a framework for spike sorting^53^ and a semi-automated spike sorting curation tool^54^, both of which were developed to be compatible with SpikeInterface. In practice, ensuring compatibility often presents significant additional work, because it involves studying other packages’ documentation, API references, and code, as well as making compromises with one’s own design choices. However, these efforts will foster a more collaborative and sustainable software landscape in neuroscience.

Neuroline achieves such compatibility by supporting different input formats for each modality and by enabling easy extensibility for new formats, tools, and modalities. Furthermore, by exporting data in the standard NWB format, Neuroline enables widely used downstream tools, such as Pynapple^32^, to read and analyse the resulting files.

## Conclusion

Contemporary neurophysiology research demands extensive, cross-modal analyses while producing data that can be shared according to FAIR principles. Neuroline aims to make a practical step forwards by modular integration of different software packages, handling metadata and format conversions between them, and exporting the combined data in the NWB format. Future development should focus on broader modality coverage, dependency and update sustainability, and compatibility and validation across datasets and laboratories.

## Methods

### Software & packages

Neuroline was developed with Python 3 and depends heavily on the NumPy^55^, SciPy^56^, pandas^57^, and Matplotlib^58^ packages. To process electrophysiological data, the packages SpikeInterface^24^, Open Ephys Python Tools^33^, and Phy^59^ are used. Suite2p^34^ is used for processing two-photon imaging data; OpenCV^60^ for video processing; DeepLabCut^41^ with the PyTorch^61^ backend for pupil extraction from video data; and FaceMap^42^ for facial motion estimation. To store the processed data in an NWB file, we use the PyNWB^31^ and NeuroConv^48^ packages.

### Electrophysiology use case

#### Experimental data

For a demonstration of the functionality of the pipeline, we used an extracellular electrophysiological recording from a 6-month-old male mouse (C57BL/6, Envigo) trained to perform a multisensory change-detection task. Here we give a brief overview of the experimental procedures. For full details of the task and similar involved procedures we refer to a previously published article^5^. All experiments were approved by the Dutch Committee for Animal Experiments and by the local Animal Welfare body of the University of Amsterdam.

To allow for head-fixation during training and data collection a titanium head-bar was surgically attached to the skull of the animal under anaesthesia (isoflurane; 4-5% for induction, ∼2% for maintenance) at the start of the experiment. After a period of recovery and habituation to head-fixation (1-2 weeks), animals were first trained to report changes in both unimodal conditions (auditory-only, visual-only) with gradually increasing saliencies, before transitioning to the full multimodal change-detection task. Stimuli were continuously presented to the animal and consisted of weighted combinations of pure tones ranging from 2^13^ Hz (=8,192 Hz) to 2^14^ Hz (=16,384 Hz) delivered through two tweeters positioned on either side of the mouse (approximately 10 cm from the ears); or full-screen drifting square-wave gratings presented on a monitor (60 Hz refresh rate) positioned 20 cm in front of the mouse. Visual gratings were presented at 70% contrast and had a temporal frequency of 1.5 Hz and a spatial frequency of 0.08 cycles per degree. A change in the auditory modality involved an instantaneous change in frequency of the auditory stimulus, a change in the visual modality involved a change in orientation of the visual grating. The animal was rewarded when licking to the correct lickspout (i.e. right for auditory changes, left for visual changes). Training animals for this task took about 8-12 weeks.

Before the extracellular recording, a small craniotomy (∼1 mm) was made over the auditory cortex under anaesthesia based on stereotactic coordinates (relative to bregma: −2.6 AP, 4.3 ML). The extracellular recording was made in the primary auditory cortex (A1) using a high-density silicon probe (A1x64-Poly2-6mm-23s-160-A64, NeuroNexus, Ann Arbor, USA). The analogue signal was preamplified and bandpass filtered (0.1–20 kHz) via an Intan RHD2164 amplifier (Intan Technologies) and continuously sampled at 30 kHz using the Open Ephys Acquisition Board (3rd generation) running the Open Ephys GUI.

Orofacial movements were captured with a monochrome camera (acA1300-200um, Basler, Ahrensburg, Germany) fitted with a 50 mm f/2.8 zoom lens (Navitar), mounted approximately 20 cm from the left side of the animal. Infrared illumination (850 nm LED) provided high-contrast imaging of the face. A video of the face of the mouse was acquired at 100 fps at a resolution of 1280 × 1024 pixels. Task-related events (e.g. ‘stimulus-change’ times), behaviour-related events (e.g. lick times) and camera acquisition events (e.g. exposure active times) were sent as digital inputs to the Open Ephys Acquisition board, either using an Arduino (Arduino Uno Rev3) or directly using a line output of the camera.

#### Processing pipeline

*EphysStream.* The raw electrophysiological data were preprocessed and spike sorted using a single call to the *run_spikesorting()* method (see *Code Availability* for use case scripts). For this recording, we processed the data in three steps. First, the data were high-pass filtered with a cutoff frequency of 400 Hz. Next, noisy or dead recording channels (e.g. channels with abnormally high or low amplitude, respectively) were automatically detected by looking at the coherence and power spectral density (PSD) between neighbouring channels, a method introduced by the International Brain Laboratory^62^. These recording channels were then interpolated using adjacent channels. Finally, to reduce common noise artefacts, the median of the raw traces of nearby channels (excluding the two directly neighbouring channels) was subtracted from the high-pass-filtered data. Spike sorting was then performed by the *MountainSort5* algorithm^63^ via SpikeInterface. Next, the sorting was analysed using Neuroline’s *analyse_sorting()* method, and several commonly reported unit quality metrics^64^ were automatically calculated (e.g. the number of inter-spike-interval (ISI) violations). Automatically detected clusters were then manually curated using Phy^59^. Clusters were classified as putative good ‘single units’ based on the shape of the mean waveform, the amount of ISI-violations, and a stable presence of the cluster throughout the session. *BehaviourStream.* The raw video data was used for both the calculation of the estimated facial motion (*FacialMotion*) and the extraction of the pupil size (*TrackedPupil*). The facial motion energy was automatically computed by Neuroline and used Facemap’s implementation^42^. The absolute difference between pixel intensities of consecutive frames was calculated, and singular value decomposition (SVD) was applied to this difference to obtain a total motion energy per frame, defined as the sum of absolute values across all principal components. The obtained motion trace was further preprocessed by low-pass filtering (cut-off frequency: 10 Hz). *TrackedPupil* was automatically computed within Neuroline by detecting 16 key points (8 on the pupil edge and 8 on the eye) in each video frame using a pre-trained DeepLabCut model. Next, an ellipse was fitted to the pupil and eye key points separately using the direct least-squares method^65^. Blinks were automatically detected and linearly interpolated using the cubic-spline method. The pupil size trace was further preprocessed by low-pass filtering (cut-off frequency: 10 Hz) and normalisation (dividing the signal by the 99.5^th^ percentile). These pupil-area preprocessing steps were inspired by Mathôt et al (2018)^66^.

#### Generalized Linear Model

The created NWB file was loaded using the Pynapple package^32^, and peri-event time histograms (PETHs) were made by aligning every individual stream to the stimulus-change event TTL. Next, we fit a generalized linear model (GLM) using the NeMoS package^30^ (see *Code Availability*). The model was trained to predict the firing rate of individual neurons in 10 ms bins over a window spanning 1 s before to 2 s after stimulus change. We included three sets of predictors in the model: a ‘stimulus’ predictor set (i.e. stimulus change TTL time), a facial motion predictor and a pupil size predictor. For the stimulus predictor set, we used a linearly-spaced raised cosine basis function with eight basis functions and a window size of 300 ms. All predictor sets were z-scored before model fitting. In total, we fit four individual models per neuron: three models that included only their respective predictor set, and a full model including all three sets of predictors. Individual models were fitted using proximal gradient descent and used an elastic net regularizer (L1 + L2 regularisation, regularizer strength: 1×10^−7^). Additionally, we used five-fold cross-validation.

#### Two-photon imaging use case

The two-photon imaging data were obtained from a recording of spontaneous activity taken from Bassler et al. (2026)^40^, which can be referred to for more details considering the experiments. These experiments were approved by the Dutch Committee for Animal Experiments and by the Animal Welfare Body of the University of Amsterdam. Recordings were performed in 8 to 52-weeks old double transgenic mice (Rasgrf2-2A-dCre;Ai148D) expressing the GCaMP6f calcium activity reporter. A head bar was surgically attached to the mouse’s skull for head-fixation under anaesthesia (4-5% isoflurane for induction; 1-2% isoflurane for maintenance). Next, a craniotomy with a diameter of 4 mm was made over the left visual cortex and subsequently, a glass cranial window was implanted. After surgery, the mice received carprofen and enrofloxacin in their drinking water for at least three days.

A near-infrared laser (Spectra-Physics Mai Tai Ti:Sapphire, MKS Instruments, Massachusetts, USA; 920 nm) was used to excite GCaMP6f. A Leica DM6000B (Leica Camera AG, Germany) microscope combined with a 16x objective (Leica HC FLUOTAR L 16x/0,60 IMM CORR VISIR) was used to image a 570×570 μm plane at a depth of 100-200 μm with a resolution of 512×512 pixels using a resonant mirror scanner operating at 27.5 Hz. Awake or anaesthetised animals were head-fixed to a custom head bar holder placed on the microscope stage.

The TIFF file consisting of the recording was loaded into Neuroline using a *TwoPhotonStream* object, which automatically generated a *TwoPhotonRecording* instance. Using this recording, Suite2p^34^ was run, yielding a *ProcessedTwoPhotonRecording* object. On this object, the *to_nwb()* method was called to add the data to the NWB file, specifically, the raw fluorescence, deconvolved fluorescence obtained using Suite2p’s implementation of the OASIS algorithm^67^, and neuropil signals; the plane segmentation; and the mean image. For visualisation, the raw cell fluorescence was neuropil-corrected by subtracting 70% of the neuropil signal, after which ΔF/F_0_ was computed using the tenth percentile of each trace as F_0_. Finally, each trace was filtered using a one-dimensional Gaussian kernel (σ = 75 ms).

## Data Availability

A dataset used to run Neuroline’s test suite as well as the raw and processed data for the use cases are available on Figshare (https://doi.org/10.21942/uva.33368623)^68^.

## Code Availability

The code and documentation are available on GitLab (https://gitlab.com/csnlab/neuroline) and archived on Zenodo (https://doi.org/10.5281/zenodo.22207352)^69^ and are licensed under the free software licence GPLv3. Example and use case scripts used in this paper are also available online (https://gitlab.com/csnlab/andreasknoben/neuroline-paper-resources).

## Author Contributions

A.K.: conceptualisation, methodology, software, validation, formal analysis, data curation, writing - original draft, writing - review & editing, visualisation. A.F.: conceptualisation, methodology, software, validation, formal analysis, data curation, writing - original draft, writing - review & editing, visualisation. S.K.: methodology, software, validation, investigation, data curation, writing - original draft, writing - review & editing, visualisation. M.B.: investigation, resources. C.M.A.P.: supervision, writing - review & editing, funding acquisition. C.A.B.: conceptualisation, writing - review & editing, supervision, funding acquisition. U.O.: conceptualisation, writing - review & editing, supervision, funding acquisition.

## Acknowledgements

We thank Toon Brouwer, Sabine Gnodde, Amber van Mierlo, Kengo Takahashi, and Junying Wang for early testing of and providing feedback on Neuroline as well as providing data to test Neuroline with different experimental setups.

## Ethics Statement

All experiments were approved by the Dutch Committee for Animal Experiments and by the local Animal Welfare Body of the University of Amsterdam (CCD application number AVD11100202216078).

