## Supplementary Figure 1 for "Neuroline: an integrated modular pipeline for FAIR multimodal neurophysiology data processing"

### Supplementary Materials

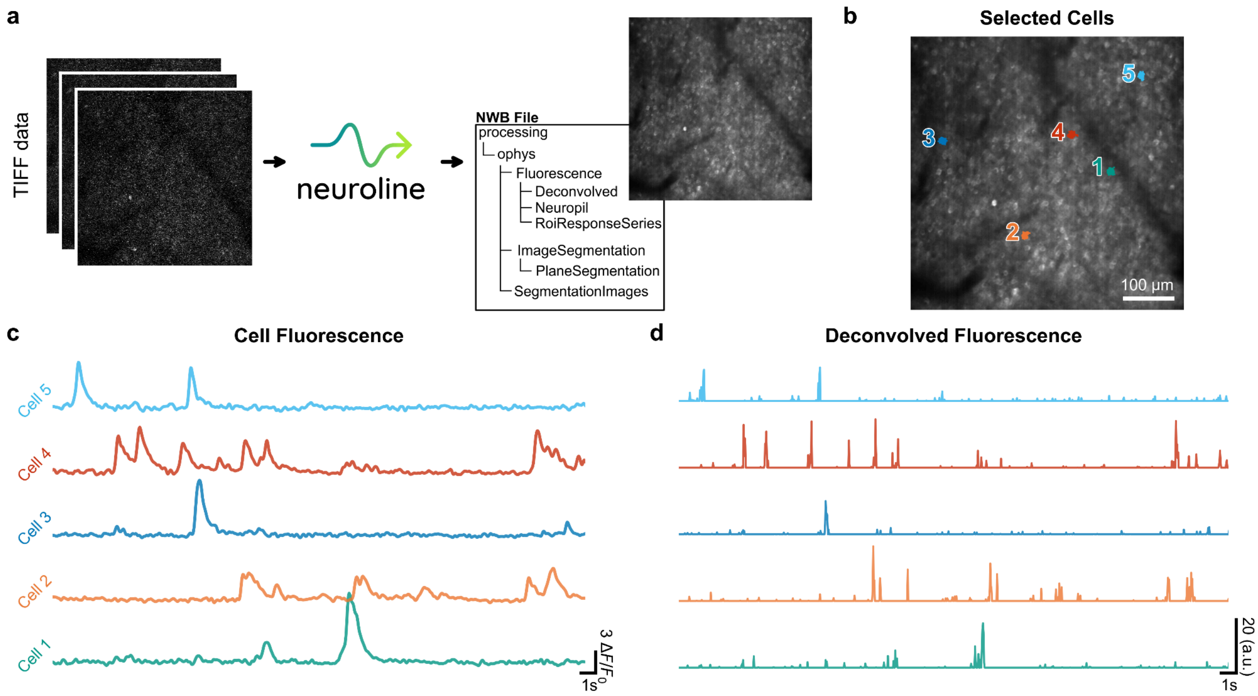


**Supplementary Figure 1. Two-photon imaging data run through Neuroline. a)** TIFF image data can be loaded into Neuroline, creating a *TwoPhotonRecording* object. Calling *run_suite2p()* runs Suite2p[1] and creates a *ProcessedTwoPhotonRecording* object. Calling *to_nwb()* on this processed data object then places the results (fluorescence traces, plane segmentations, and images) in an NWB file using NeuroConv[2]. Data used here is spontaneous activity from Bassler et al., 2026[3]. Two-photon imaging was conducted over left mouse visual cortex using the GCaMP6f calcium indicator[3]. **b)** Mean fluorescence image overlaid with the selected cells (coloured). Numbers next to each cell correspond to the numbered traces in panel c and d. **c)** Fluorescence of selected cell bodies. Raw fluorescence was corrected by subtracting 70% of the neuropil signal; this corrected fluorescence signal was then used to compute $\Delta F/F_0$, where $F_0$ is the tenth percentile of the entire trace. Finally, each trace was filtered using a one-dimensional Gaussian kernel ($\sigma$ = 75 ms). **d)** The deconvolved activity of the selected cells, obtained using Suite2p's implementation of the OASIS algorithm[4]. The deconvolved traces represent inferred spiking activity from the fluorescence data, providing a signal less affected by the slower kinetics of the calcium indicator.
